# The RNA pseudoknots in foot-and-mouth disease virus are dispensable for genome replication but essential for the production of infectious virus

**DOI:** 10.1101/2020.01.10.901801

**Authors:** Joseph C. Ward, Lidia Lasecka-Dykes, Chris Neil, Oluwapelumi Adeyemi, Sarah Gold, Niall McLean, Caroline Wright, Morgan R. Herod, David Kealy, Emma Warner, Donald P. King, Tobias J. Tuthill, David J. Rowlands, Nicola J. Stonehouse

**Affiliations:** School of Molecular and Cellular Biology, Faculty of Biological Sciences and Astbury Centre for Structural Molecular Biology, University of Leeds, Leeds, United Kingdom; Pirbright Institute, Ash Rd, Pirbright, Surrey, GU24 ONF, United Kingdom; Department of Medical Microbiology and Parasitology, Faculty of Basic Medical Sciences, College of Health Sciences, University of Ilorin, Ilorin, Nigeria

**Keywords:** picornavirus, pseudoknot, 5′ UTR, replication, foot-and-mouth disease virus

## Abstract

The positive stranded RNA genomes of picornaviruses comprise a single large open reading frame flanked by 5′ and 3′ untranslated regions (UTRs). Foot-and-mouth disease virus (FMDV) has an unusually large 5′ UTR (1.3 kb) containing five structural domains. These include the internal ribosome entry site (IRES), which facilitates initiation of translation, and the cis-acting replication element (*cre)*. Less well characterised structures are a 5′ terminal 360 nucleotide stem-loop, a variable length poly-C-tract of approximately 100-200 nucleotides and a series of two to four tandemly repeated pseudoknots (PKs). We investigated the structures of the PKs by selective 2′ hydroxyl acetylation analysed by primer extension (SHAPE) analysis and determined their contribution to genome replication by mutation and deletion experiments. SHAPE and mutation experiments confirmed the importance of the previously predicted PK structures for their function. Deletion experiments showed that although PKs are not essential for replication, they provide genomes with a competitive advantage. However, although replicons and full-length genomes lacking all PKs were replication competent, no infectious virus was rescued from genomes containing less than one PK copy. This is consistent with our earlier report describing the presence of putative packaging signals in the PK region.

## Introduction

Foot-and-mouth disease virus (FMDV) is a single stranded positive sense RNA virus of the genus *Aphthovirus* in the family *Picornaviridae*. It occurs as seven, antigenically diverse serotypes; A, O, C, Asia 1, South African Territories (SAT) 1, 2 and 3. It is the causative agent of foot-and-mouth disease (FMD), a highly contagious disease of cloven-hooved animals affecting most notably cattle, pigs, sheep and goats in addition to wild species such as the African buffalo. Disease outbreaks have serious economic implications resulting from trade restrictions, reduced productivity and the slaughter of infected and at-risk animals (1). The 2001 outbreak in the UK caused economic losses of over £8 billion to the tourism and agricultural sectors. Inactivated virus vaccines are used in countries in which FMD is endemic, but these are often strain-specific and provide little cross protection between serotypes (2). Antigenic variation together with the relatively short duration of immunity following vaccination combine to complicate control of the disease (3). In addition, the carrier state, in which asymptomatically infected animals continue to shed virus, contributes to the spread of FMDV (4). An improved understanding of the viral life cycle may be important for the development of improved vaccines and other control measures.

The FMDV genome (approximately 8.4 kb) consists of a single open reading frame flanked by 5′ and 3′ untranslated regions (UTRs) (figure 1A) (5). The translated region encodes both structural and non-structural proteins. The P1 region encodes the capsid structural proteins VP1, VP3, and VP0 (which is further processed to VP2 and VP4 during virus assembly) (6). The P2 and P3 regions encode the non-structural proteins 2B and 2C and 3A, 3B_(1-3)_ (VPg), 3C^pro^ and 3D^pol^ respectively (7, 8). Due to disease security restrictions, work with infectious virus is limited to a few high containment facilities. However, sub-genomic replicons, in which the structural protein-coding region is replaced by reporter genes, allow the study of genome replication without the requirement for high containment (9, 10) (figure 1A).

**Figure 1.**
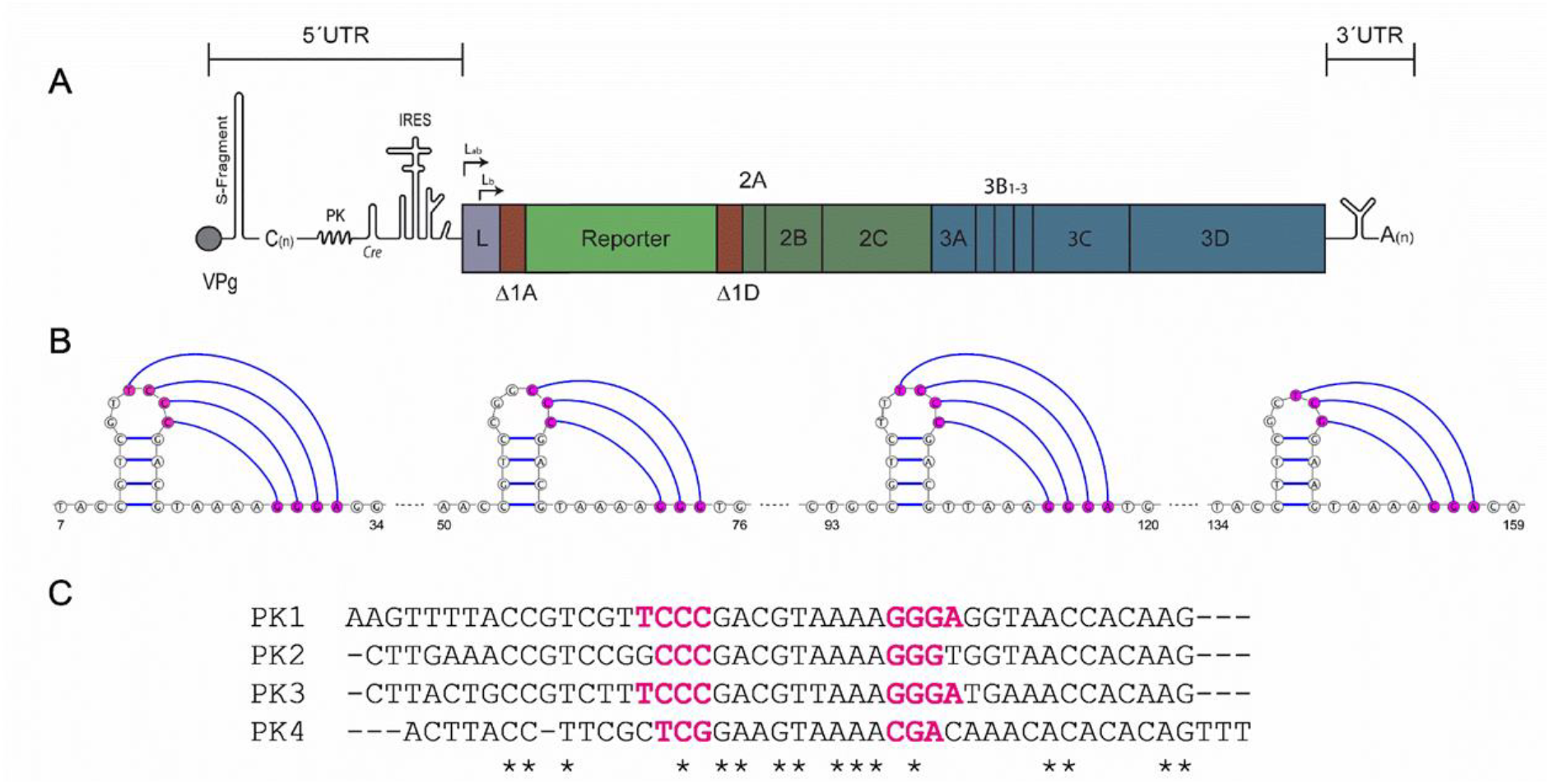
Replicon and PK schematic. Schematic of the FMDV O1K sub-genomic replicon, showing both 5’ and 3’ untranslated regions (UTRs) together with the RNA structures present in these regions. IRES-driven translation produces a single polyprotein. Here, the structural proteins have been replaced with a green fluorescent reporter, upstream of the non-structural proteins 2A-3D (A). Predicted PK structures, with putative interactions highlighted in hot-pink are shown. Numbers indicate nucleotide positions after the poly-C-tract (B). Sequence alignment of the 4 PKs, with the interacting regions shown in hot-pink and invariant nucleotides represented by asterisk (C).

The FMDV 5′ UTR is the largest known picornavirus UTR, comprising approximately 1300 nucleotides and containing several highly structured regions. The first 360 nucleotides at the 5′ end are predicted to fold into a single large stem loop termed the S-fragment, followed by a large poly-C tract of variable length (which can be up to 200 nt), a region containing two to four tandemly repeated pseudoknots (PKs), the cis acting replication element (*cre*) and the internal ribosome entry site (IRES) (5, 11, 12). Of these five structural domains, functions have been ascribed to only two, the *cre* and IRES. The *cre* region is involved in uridylation of the RNA primer peptide, VPg (also known as 3B), and the IRES determines the initiation of translation of the viral polyprotein (13, 14). The roles of the S-fragment, the poly-C tract and the PKs in viral replication are not fully elucidated, however recent studies have shown that truncations to the S-fragment can play key roles in the innate immune response to viral infection (15–17). It has also recently been reported that viruses containing a deletion within the pseudoknot region showed an attenuated phenotype in bovine cell lines while remaining unchanged in porcine, suggesting a role for the pseudoknots in viral tropism (18).

The PKs were originally predicted in 1987 and consist of two to four tandem repeats of a ~48 nucleotide region containing a small stem loop and downstream interaction site (figure 1B) (12). Due to the sequence similarity between the PKs (figure 1C), it is speculated that they were formed by duplication events during viral replication, probably involving recombination. Between two and four PKs are present in different virus isolates but no strain has been identified with less than two PKs, emphasising their potential importance in the viral life cycle (19, 20). The presence of PKs has been reported in the 5′ UTR of other picornaviruses such as encephalomyocarditis virus (EMCV) and equine rhinitis A virus (ERAV) (21, 22). However, in both cases the PKs are located at the 5′ side of the poly-C-tract, making their location in the FMDV genome unique.

PKs have been reported to have roles in several aspects of viral replication including splicing (e.g. HIV and influenza), ribosomal frameshifting (e.g. coronaviruses) and RNase protection (e.g. Dengue virus) (23–27). In the work reported here, the role of the PKs in the FMDV life cycle was investigated, together with biochemical probing of PK structures. The combination of both virus and replicon systems allowed us to distinguish effects on genome replication and other aspects of the viral life cycle. Selective mutation within the PK domain and sequential deletion of PKs confirmed the importance of PK structure and that although genome replication can occur in the absence of PKs at least one is required for wild-type (wt) replication. Furthermore, competition experiments showed that extra copies of PKs conferred a replicative advantage to genomes. Although replicons and full-length genomes lacking PKs were replication-competent, no infectious virus was rescued from genomes containing less than one PK copy. This is consistent with our earlier report describing the presence of putative packaging signals in the PK region (22).

## Materials and Methods

### Cells lines

BHK-21 cells obtained from the ATCC (LGC Standard) were maintained in Dulbecco’s modified Eagle’s Medium with glutamine (Sigma-Aldrich) supplemented with 10 % foetal calf serum (FCS), 50 U/ml penicillin and 50 µg/ml streptomycin.

### Plasmid construction

The FMDV replicon plasmids, pRep-ptGFP, and the replication-defective polymerase mutant control, 3D-GNN, have already been described (10).

To introduce mutations into the PK region, the pRep-ptGFP replicon plasmid was digested with *Spe*I and *Kpn*I and the resulting fragment inserted into a sub-cloning vector (pBluescript) to create the pBluescript PK. PKs 3 and 4 were removed by digestion with *Hind*III and *Aat*II before insertion of a synthetic DNA sequence with PK 3 and 4 deleted. PKs 2, 3 and 4 were deleted by PCR amplification using ΔPK 234 Forward primer and FMDV 1331-1311 reverse primer, the resultant product was digested with *Hind*III and *Aat*II and ligated into the pBluescript PK vector. Complete PK deletion was achieved by introduction of an *Afl*II site at the 3′ end of the poly-C tract by PCR mutagenesis to create the sub-cloning vector, pBluescript C11, which was then used to remove all the PKs by PCR mutagenesis using ΔPK 1234 forward primer and FMDV 1331-1311 reverse primer. The modified PK sequences were removed from the sub-cloning vectors and inserted into the pRep-ptGFP plasmid using *Nhe*I-HF and *Kpn*I-HF.

Mutations to disrupt and reform PK structure were introduced using synthetic DNA by digestion with *Afl*II and *Aat*II and ligation into a similarly digested pBluescript PK vector. Mutations were then introduced into the replicon plasmid as described above.

To assess the effects of truncation of the poly-C-tract on replication the entire sequence was removed. This was performed by PCR mutagenesis using primers C0 *Spe*I, and FMDV 1331-1311 as forward and reverse primers respectively. The PCR product was digested with *Spe*I and *Kpn*I before ligation into a *Nhe*I and *Kpn*I digested *wt* pRep ptGFP replicon. Sequences of all primers are available upon request.

### *In vitro* transcription

*In vitro* transcription reactions for replicon assays were performed as described previously (28). Transcription reactions to produce large amounts of RNA for SHAPE analysis were performed with purified linear DNA as described above, and 1 μg of linearised DNA was then used in a HiScribe T7 synthesis kit (NEB), before DNase treatment and purification using a PureLink RNA mini kit (Thermo Fisher).

### SHAPE analysis

RNA was prepared as above and a sample (12 pmol) was heated to 95°C for 2 minutes before cooling on ice. RNA folding buffer (100 mM HEPES, 66 mM MgCl_2_ and 100 mM NaCl) and RNase Out (Invitrogen) was added to the RNA and incubated at 37°C for 30 minutes. Once folded, RNA was treated with NMIA compound at a final concentration of 5 mM or DMSO as a negative control for 50 minutes at 37°C. Following incubation, labelled RNA was ethanol precipitated and resuspended in 10 μl 0.5 x TE buffer.

Primer extension of NMIA labelled RNA was performed by incubation of 5 μl of labelled RNA with 6 μl of RNase free water and 1 μl of 10 μM Hex of FAM fluorescent primer. Primer binding was facilitated by heating the reaction to 85°C for 1 minute, 60°C for 10 minutes and 35°C for 10 minutes in a thermocycler. A reverse transcription master mix containing 4 μl first strand buffer, 1 μl 100 mM DTT, 0.5 μl RNase Out, 1 μl Supsercript III (Invitrogen), 1 μl 10 mM PCR dNTP mix (Promega) and 0.5 μl RNase free water, was then added to the RNA/primer complex and extension carried out by incubation at 52°C for 30 minutes.

Post extension, cDNA:RNA hybrids were disassociated by incubation with 1 μl 4M NaOH at 95°C for 3 minutes before neutralisation with 2 μl 2M HCl. Extended cDNA was ethanol precipitated and resuspended in 40 μl deionized formamide (Thermo Fisher). Sequencing ladders were made similarly using 6 pmol of RNA with the inclusion of 1 μl 10 mM ddCTP in the reverse transcription mix and using a differentially labelled fluorescent primer (either Hex or FAM). 20 μl of sequencing ladder was combined with NMIA or DMSO samples and dispatched on dry ice for capillary electrophoresis (Dundee DNA seq).

Capillary electrophoresis was analysed using QuShape and reactivity overlaid onto the RNA structure using VARNA (29, 30).

### Replication assays

Replicon replication in all cell lines was assessed in 24-well plates with 0.5 µg/cm^2^ of RNA using Lipofectin transfection reagent (Life Technologies) as previously described (28). For complementation assays, BHK-21 cells seeded into 24-well plates were allowed to adhere for 16 hours before transfection with 1 µg of replicon RNA using Lipofectin. Each transfection was performed in duplicate and experiments were biologically repeated. Replicon replication was assessed by live cell imaging using an IncuCyte Zoom Dual colour FLR, an automated phase-contrast and fluorescence microscope within a humidifying incubator. At hourly intervals up to 24 hours post transfection, images of each well were taken and used to count the number of ptGFP positive cells per well.

Passaging in competition assays was performed by co-transfecting BHK-21 cells with *in vitro* transcribed replicon RNA and harvesting total cell RNA at 8 hours post transfection using TRIzol reagent (Thermo Fisher Scientific). The harvested RNA was then purified using the Direct-zol RNA MiniPrep kit (Zymo Research) with on-column DNase I treatment and eluted in DEPC treated water. The purified passaged RNA (1 µg) was transfected onto the naïve BHK-21 cells as above.

### Construction of recombinant viruses

Replicons used here are based on plasmid T7S3 which encodes a full length infectious copy of FMDV O1 Kaufbeuren (31). The reporter was removed from replicons by digestion with *Psi*I and *Xma*I restriction enzymes and replaced with the corresponding fragment from pT7S3 encoding the capsid proteins. Full length viral RNA was transcribed using a T7 MEGAscript kit (Thermo Fisher Scientific), DNase treated using TurboDNase (Thermo Fisher Scientific) and purified using a MEGAclear Transcription Clean-Up kit (Thermo Fisher Scientific). RNA quality and concentration were determined by denaturing agarose gel electrophoresis and Qubit RNA BR Assay Kit (Thermo Fisher Scientific).

### Virus recovery

BHK-21 cells were transfected in 25 cm^2^ flasks with 8 µg per flask of infectious clone-derived RNA using TransIT transfection reagent (Mirus) as described previously (32). 24 hours post-transfection cell lysates were freeze-thawed and clarified by centrifugation. Clarified lysate was blind passaged onto naïve BHK-21 cells, this was continued for five rounds of passaging.

### Sequencing of recovered virus

Recovered viruses at passage 4, were sequenced using an Illumina Miseq (illumine) using a modified version of a previously described PCR-free protocol ((32, 33)). Total RNA was extracted from clarified passage 4 lysates using TRizol reagent (Thermo Fisher Scientific) and residual genomic DNA removed using DNA-free DNA removal Kit (Thermo Fisher Scientific). RNA was precipitated using 3 M sodium acetate and ethanol, 10 ul of purified RNA (containing 1 pg to 5 µg) of RNA was used in a reverse transcription reaction as previously described (33, 34). Following reverse transcription cDNA was purified and quantified using a Qubit ds DNA HS Assay kit (Thermo Fisher Scientific) and a cDNA library prepared using Nextera XT DNA Sample Preparation Kit (Illumina). Sequencing was carried out on the MiSeq platform using MiSeq Reagent Kit v2 (300 cycles) chemistry (Illumina).

FastQ files were quality checked using FastQC with poor quality reads filtered using the Sickle algorithm. Host cell reads were removed using FastQ Screen algorithm and FMDV reads assembled *de novo* into contigs using IDBA-UD (35). Contigs that matched the FMDV library (identified using Basic Local ALighnment Search Tool (BLAST)) were assembled into consensus sequences using SeqMan Pro software in the DNA STAR Lasergene 13 package (DNA STAR) (36).

### Plaque assay of recovered virus

Confluent BHK-21 cell monolayers were infected with 10-fold serial dilutions of virus stock, overlaid with Eagle overlay media supplemented with 5 % tryptose phosphate broth solution (Sigma Aldrich), penicillin (100 units/ml and streptomycin (100 µg/ml) (Sigma Aldrich) and 0.6 % Indubiose (MP Biomedicals) and incubated for 48 hours at 37°C. Cells were fixed and stained with 1 % (w/v) methylene blue in 10 % (v/v) ethanol and 4 % formaldehyde in PBS.

Fixed plaques were scanned and images measured using a GNU Image Manipulation Program IMP (GIMP, available at https://www.gimp.org). For each plaque, horizontal and vertical diameter in pixels was taken and an average of these two values was calculated. All plaques per well were measured.

### Cell killing assays

Virus titre was determined by plaque assays. BHK-21 cells were seeded with 3 x10^4^ cells/well in 96 well plates and allowed to settle overnight. Cell monolayers were inoculated with each rescued virus at MOI of 0.01 PFU for 1 hour, inoculum was removed and 150 µl of fresh GMEM (supplemented with 1 % FCS) was added to each well. Appearance of CPE was monitored every 30 minutes using the IncuCyte S3.

### Flow cytometry assay

Production of viral proteins in BHK-21 cells transfected with infectious copy transcripts and Lipofectamine 2000 was measured using immunofluorescence with 2C2, a mouse antibody recognising viral protein 3A, and a goat anti-Mouse IgG (H+L) highly cross-adsorbed secondary antibody, Alexa Fluor 488 (Life Technologies). Each transcript was transfected in triplicate and the experiment biologically repeated three times. BHK-21 cells were seeded into T25 flasks 16 hours prior to transfection with 10 µg RNA. The transfection mix was left on the cells for 1 hour before the media was changed to VGM (Glasgow Minimum Essential Medium (Sigma-Aldrich), 1% Foetal Bovine Serum – Brazil origin (Life Science Production) and 5% Tryptose Phosphate Broth (Sigma-Aldrich).

After a further 3 hours, cells were dissociated using trypsin-EDTA 0.05% phenol red (Life Technologies), pelleted at 200 g for 3 minutes and fixed in 4% paraformaldehyde for 40 minutes. Cells were then transferred to a 96-well u-bottom plate and pelleted; this and all subsequent pelleting steps were done at 300 xg for 5 minutes. Cells were resuspended in 0.5% BSA in PBS blocking buffer (Melford), pelleted and resuspended in 1/1000 2C2 antibody and left shaking at 500 rpm at 4°C for 14 hours in an Eppendorf Thermomixer C plate shaker. The cells were pelleted and subsequently resuspended in blocking buffer three times to wash, resuspended in 1/200 anti-mouse fluorescent secondary antibody and rotated at 500 rpm at 24°C for 1 hour before washing a final three times. Cells were then resuspended in 500 µl PBS and data were collected on the LSR Fortessa (BD Biosciences) using BD FACSDivaTM software. Data were exported as flow cytometry standard (FCS) files, and were analysed in FlowJo 10 using the gating strategy shown in Figure 7.

## Results

### SHAPE analysis of PK structure

The presence of PKs was initially predicted in 1987 by computational and visual analysis of the 5′ UTR sequence (12). The prediction of the presence of multiple PKs was strengthened by the observation that variation in the length of this region between different virus isolates equated to the gain or loss of PK-length sequence units. However, the definitive demonstration of PK structure remains a challenge. Here, we used selective 2′ hydroxyl acylation analysed by primer extension (SHAPE) to investigate the secondary structure of the PK region.

RNA transcripts representing FMDV UTRs were folded prior to treatment with NMIA, a compound that forms 2′-O-adducts when interacting with non-paired nucleotides, or DMSO as a negative control. Labelled RNAs were purified and used as templates in reverse transcription reactions using fluorescently labelled primers. Elongation of the reverse transcription products terminates at adducts, resulting in cDNA fragments of different lengths, which were analysed by gel electrophoresis alongside a sequencing ladder to identify sites of NMIA interaction. The whole PK region was surprisingly reactive suggesting that it was largely single stranded or highly flexible (figure 2A). To investigate if the SHAPE data agreed with the predicted structure, the NMIA reactivity was overlaid onto the previous PK structure prediction (12).

**Figure 2.**
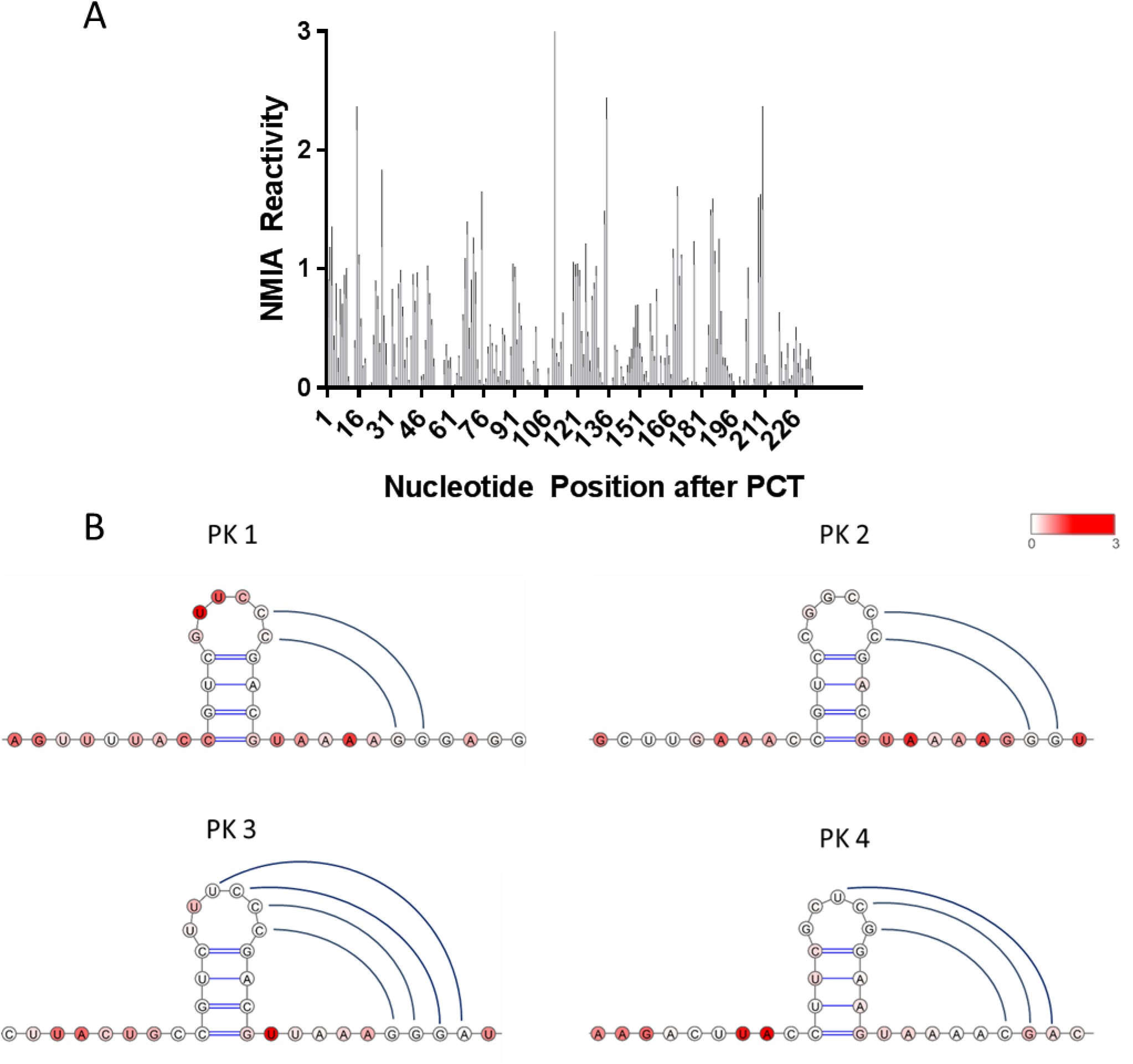
SHAPE NMIA reactivity of the PK region. NMIA reactivity at nucleotides following the poly-C-tract (PCT). High reactivity indicates increased chance of the nucleotide being non base-paired at that position (A). NMIA reactivity of each PK overlaid onto the predicted PK structure using VARNA (30). Loop and downstream interactions represent those supported by SHAPE data (B). NMIA reactivity is represented on a colour scale from low (white) to high (red) (n = 4).

The SHAPE data largely agreed with the predicted structures with the stems of PK 1, 2 and 3, being unreactive, suggestive of base-pairing. Formation of the stem of PK4 was less convincing, although the stem nucleotides still had relatively low reactivity in agreement with the other PK models. For all the PKs, the nucleotides in the loop regions and the predicted downstream interacting nucleotides showed little to no reactivity, suggesting NMIA could not interact with these nucleotides either due to the predicted base pairing or steric hindrance (figure 2B). The NMIA reactivity for the interacting nucleotides in the stem-loops with downstream residues of PK 1, 2 and 3 again largely agreed with the predicted structure, although the SHAPE data suggests that there might be fewer interactions than previously predicted. However, differences here could be due to heterogeneity in the formation of PKs in this experiment. The evidence for loop-downstream interaction was weaker for PK4.

By overlaying the NMIA reactivities with the original predicted structure the SHAPE data were compatible to the PK models and potentially shed new light on the requirements of the loop interactions.

### A single PK is sufficient for efficient replication

The replicon system was based on the O1K FMDV sequence which includes four similar but non-identical PKs (figure 1). The PKs were sequentially deleted from the 3′ side (i.e. PK 4-PK 1), and replication of the resulting modified replicons assessed.

To allow complete removal of all PKs, an *Afl*II site was inserted into the ptGFP replicon plasmid which resulted in reduction of the poly-C-tract to 11 cytosine residues. This C11 replicon was investigated alongside a *wt* replicon and one with lethal polymerase mutations (3D-GNN). These controls were used to confirm that truncation of the poly-C tract had no measurable effect on replication the two cell lines tested, as previously reported (37) (figure 3A). For completeness, we further removed the entire poly-C-tract (C0) and showed that this had no observable negative effect on replication of the replicon (figure 3B). The C11 construct was then used as the “backbone” for removal of all four PKs.

**Figure 3.**
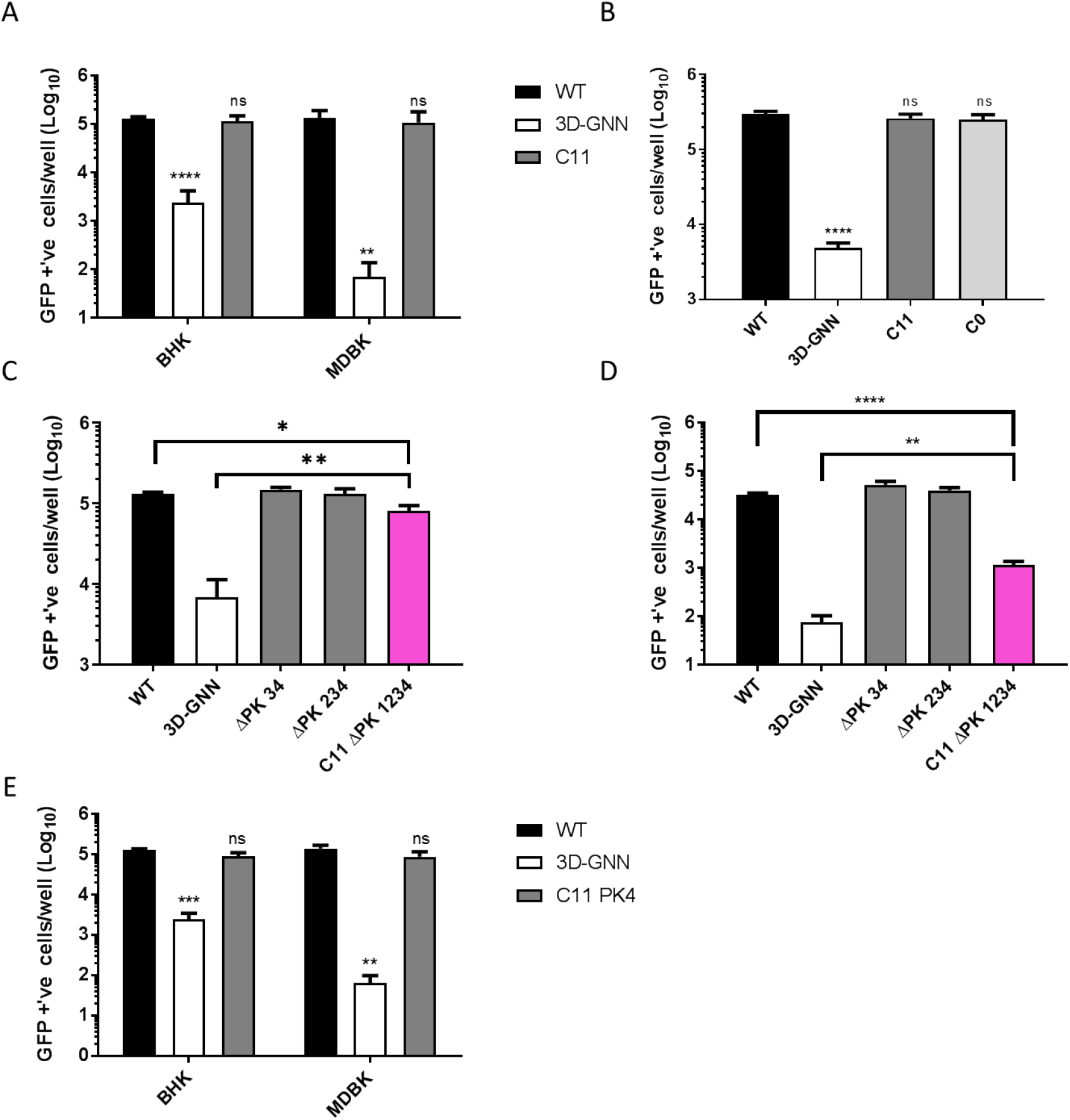
The poly-C-tract is dispensable and only one PK is required for *wt* replication. *Wt*, 3D-GNN and a replicon with a truncated poly-C-tract (C11) were transfected into BHK-21 and MDBK cell lines, replication was monitored using the Incucyte Zoom (A). A replicon with entire poly-C-tract removed (C0) was transfected alongside *wt*, 3D-GNN and C11 replicons into BHK-21 cells (B). Replicons with sequentially deleted PKs (ΔPK 34, ΔPK 234 and C11 ΔPK 1234) were assayed for replication in BHK-21 cells (C) and MDBK cells (D). Replication of replicon with PK 4 as the sole remaining PK (C11 PK 4), transfected into MDBK and BHK-21 cells (E). All replication assays were measured by counting the number of GFP positive cells per well using the Incucyte Zoom with data shown at 8 hours post-transection. Error bars shown are calculated by SEM, n = 3. Significance is shown compared to the *wt* (A, B) or the *wt* and 3D-GNN (C, D, E) * P < 0.05, ** P < 0.01, *** P < 0.001, **** P < 0.0001.

Replicons with PK deletions were transfected into baby-hamster kidney-21 cells (BHK-21) (figure 3C) or Madin Darby bovine kidney cells (MDBK) (figure 3D). Replication was monitored by measuring ptGFP reporter expression, in parallel with transfection of a *wt* and 3D-GNN replicon, where the 3D-GNN replicon is used to monitor ptGFP expression resulting from translation of input RNA in the absence of replication. Reporter expression was recorded using an IncuCyte Zoom automatic fluorescent microscope and is shown at 8 hours post-transfection, analysed by our standard methods (8, 10, 38).

Following transfection into each cell line, replicons lacking PK 3, 4 and 2, 3, 4 (termed ∆PK 34 and ∆PK 234 respectively) replicated at similar levels to the *wt* replicon (figure 3C-D). However, a replicon containing no PKs (C11 ∆PK 1234) showed a significant (~ 4 fold) reduction in replication in BHK-21 cells compared to the *wt* C11 replicon. A larger reduction in replication (28 fold) was seen in the MDBK cell line, supporting previous publications on the potential role in host cell tropism (18). Replication of the C11 ∆PK 1234 replicon in MDBK cells was however still significantly above that of the 3D-GNN negative control. These data suggest that although the PKs are not essential for replication at least one PK is required for *wt* levels of replication.

In the experiments above PK1 was the sole remaining PK and we therefore investigated whether other PKs could similarly support *wt* replication. We deleted all the PKs to create the C11 construct and re-inserted PK4 as the only PK (C11 PK4). Near *wt* levels of replication were observed following transfection into both cell types suggesting that there is no functional difference between PK1 and PK4 (figure 3E).

### Function of the PKs in replication is dependent on downstream interactions and orientation

Since removal of all four PKs resulted in a significant decrease in replication, the minimal requirements to maintain *wt* levels of replication were investigated. As near *wt* level of replication was observed when only one PK was present, all further mutagenesis was performed in a C11 replicon plasmid containing only PK 1.

Mutations designed to interrupt base pairing and abrogate formation of the PK structure were made in the loop of PK 1 and the corresponding downstream nucleotides. The substitutions (shown in red) created a GAGA motif both in the loop and downstream regions and reduced the replication of the mutated replicon (C11 PK disrupt) equivalent to that of the replicon containing no PKs, thereby supporting the predicted structure (figure 4A). Base pairing potential was then restored by mutation of the relevant nucleotides in the loop and downstream region to GGGG and CCCC respectively. Restoring the interaction using an alternate sequence increased replication significantly compared to the disrupted PK replicon (~ 4 fold), although this was still slightly below that of the *wt* (~ 0.7 fold decrease) (figure 4A).

**Figure 4.**
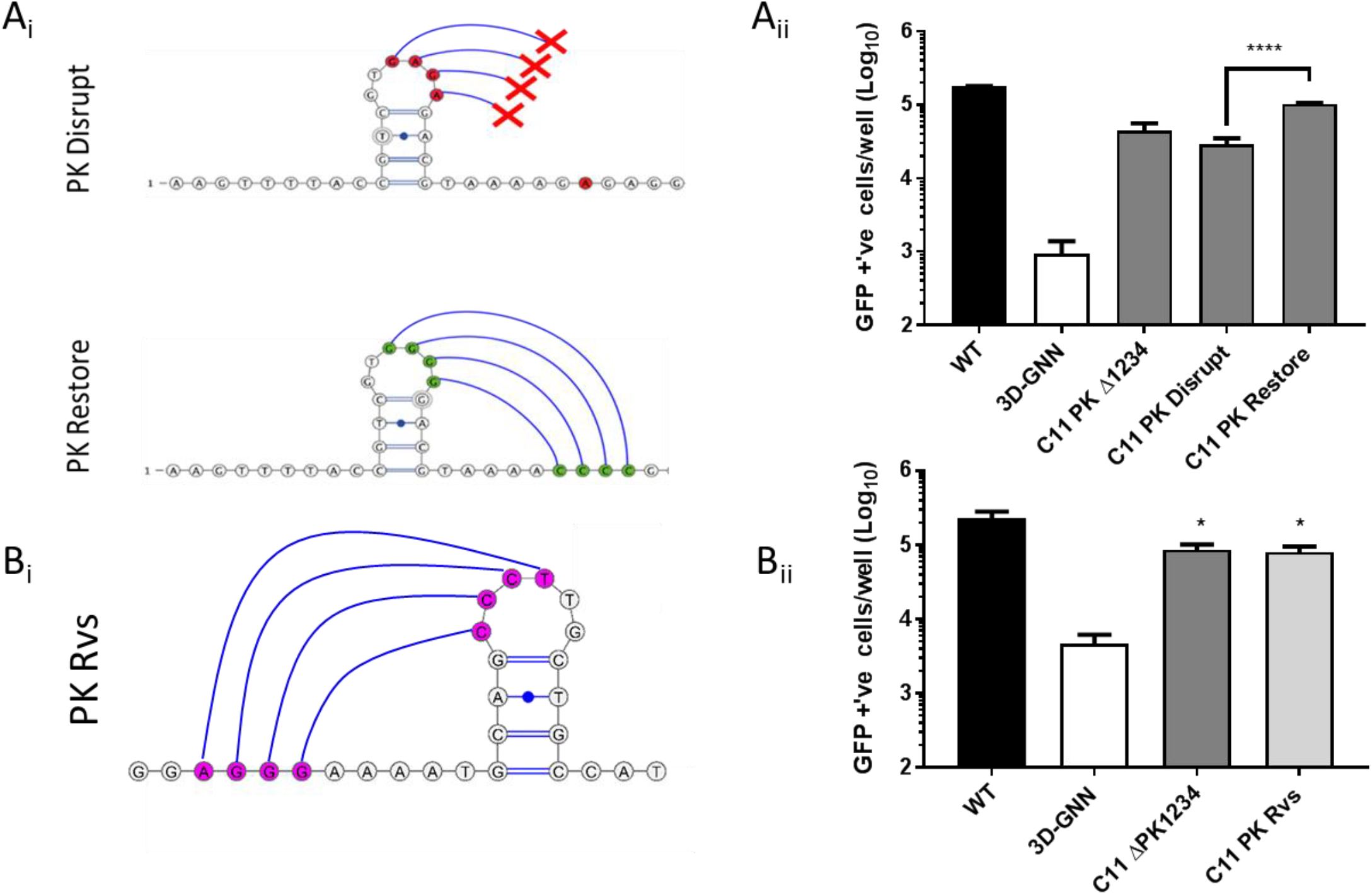
Disrupting the PK structure and reversing the orientation of a PK reduces replication. Cartoon representations of disrupting and restoring mutations made to PK 1, where nucleotides in the bulge of the stem loop and interacting region downstream were mutated to disrupt structure formation ‘PK disrupt’, or mutated to maintain bulge and downstream interaction but with different nucleotides ‘PK restore’ (A_i_). Replication of PK disrupt and restore mutants were measured by transfection of RNA into BHK-21 cells and shown here at 8 hours post-transfection alongside *wt*, 3D-GNN and C11 ΔPK 1234 controls. Significance is shown comparing the replication of C11 PK disrupt and C11 PK restore (A_ii_). Visual representation of the reversing of the nucleotide sequence of PK1 creating the C11 PK Rvs construct (B_i_). Replication of PK Rvs at 8 hours post transfection of BHK-21 cells (B_ii_). Significance shown is compared to *wt* replicon. Error bars are calculated by SEM, n = 3, * P < 0.05, **** P < 0.0001.

In addition, the orientation of PK 1 was reversed by “flipping” the nucleotide sequence to potentially facilitate hybridisation of the loop with upstream rather than downstream sequences. Changing the orientation of the PK reduced replicon replication to a similar level seen in the absence of PKs (figure 4B). This suggests that the role of the PKs in genome replication is dependent on both sequence, structure and orientation.

### Multiple PKs confer a competitive replicative advantage

The deletion studies above suggested that removal of up to three of the four PKs present in the *wt* sequence had no clear effect on replicon replication, although deletion of all four was significantly detrimental. To investigate whether multiple PKs conferred more subtle advantages for replication than were evident from single round transfection experiments we carried out sequential passages of replicon RNA following transfection of the PK deleted forms in competition with a *wt* replicon. Different reporter genes (ptGFP or mCherry) were used to distinguish the competing replicons.

Replicons encoding ptGFP; *wt*, ∆PK 34, ∆PK 234 and C11 ∆PK 1234 were co-transfected into BHK-21 cells together with either a *wt* mCherry replicon or yeast tRNA as a control. The replication of each of the co-transfected replicons was compared by observing ptGFP and mCherry expression over three sequential passages. Passaging was achieved by harvesting total RNA using Trizol-reagent 8 hours post-transfection. Harvested RNA was purified and then re-transfected into naïve BHK-21 cells.

Co-transfection of the *wt*, ∆PK 34 or ∆PK 234 with yeast tRNA as controls showed no differences in replication as expected (Figure 5A). Likewise, when PK mutants were co-transfected with a *wt* replicon after three passages, the number of green fluorescent cells produced by the ∆PK 34 replicon was comparable to that of the *wt*, suggesting no competitive advantage of four PKs over two. For both, there was a reduction in replication after the first passage but recovery to near that of the original transfection by the third passage. However, when co-transfected with the *wt* replicon, the ∆PK 234 replicon showed a similar drop in replication in passage two, but showed no subsequent recovery following each passage and replication decreased until at passage three there is a 2.5 fold reduction compared to that of passage 0 (figure 5B). Therefore, it appears that replicons with a single PK are at a competitive disadvantage compared to those with two or more.

**Figure 5.**
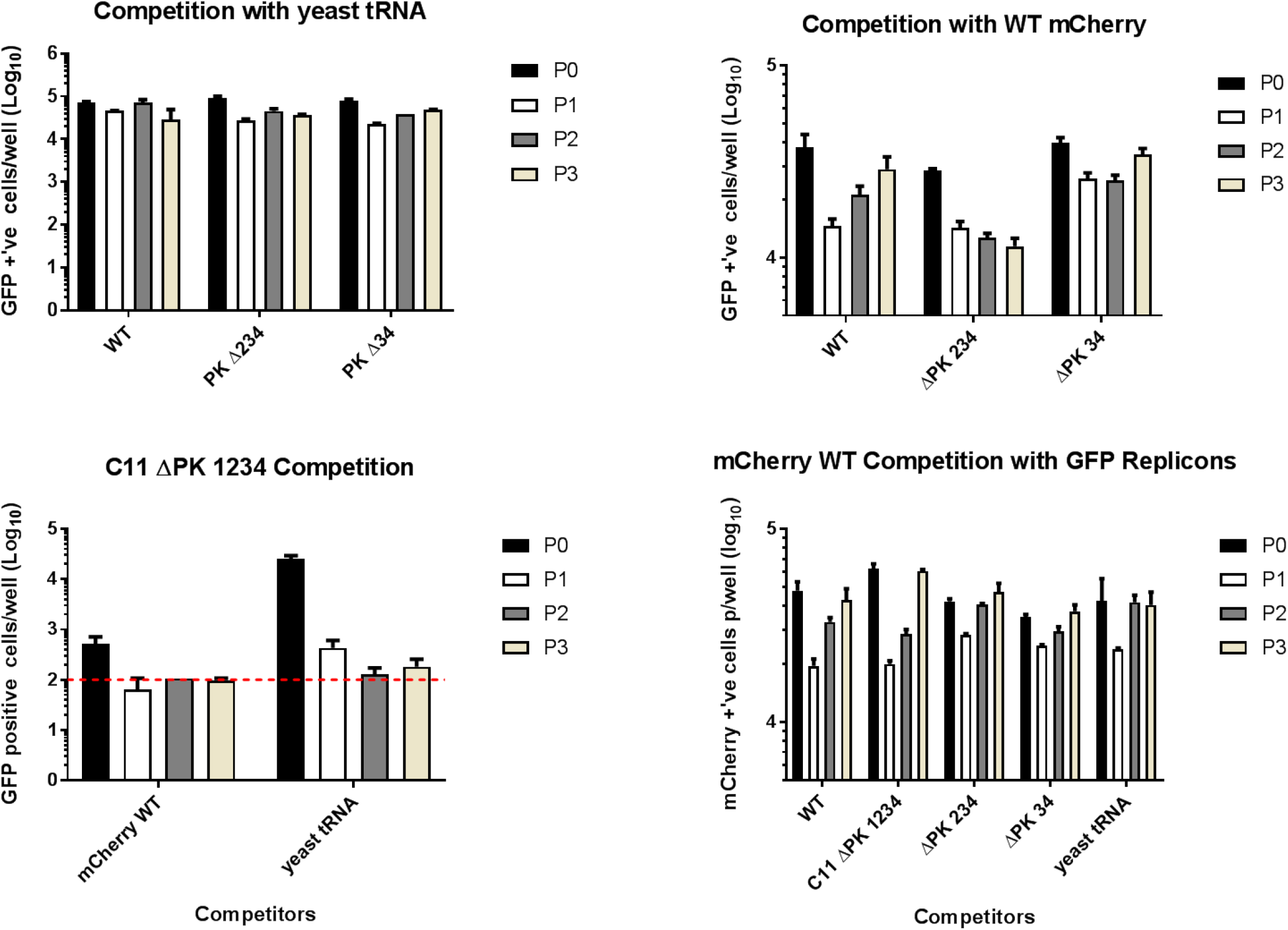
More than 2 PKs provides a replicative advantage in co-transfection competition experiments. BHK-21 cells were co-transfected with *wt*, ΔPK 1234, ΔPK 234 or ΔPK 34 ptGFP replicon RNA and either a *wt* mCherry replicon or yeast tRNA control. Control competition of ptGFP *wt*, ΔPK 234 or ΔPK 34 replicons with yeast tRNA (A). Competition of *wt*, ΔPK 234 or ΔPK 34 ptGFP replicons with a *wt* mCherry replicon (B). Replication of the C11 ΔPK 1234 replicon when co-transfected with either the *wt* mCherry replicon or yeast tRNA control (C). Replication of the *wt* mCherry replicon when co-transfected with ptGFP *wt*, C11 ΔPK 1234, ΔPK 234 or ΔPK 34 ptGFP replicons (D). All replication is shown at 8 hours post transfection over 3 sequential passages, as measured by an Incucyte Zoom (n = 2). Initial transfection (P0), sequential passages (P1-3).

Co-transfection with the *wt* mCherry replicon reduced the replication of the C11 ∆PK 1234 replicon to background levels as seen when comparing to the yeast tRNA control. By passage two the ptGFP signal of the C11 ∆PK 1234 was no longer detectable, suggesting that this replicon has been out competed (figure 5C). Although the initial replication of C11 ∆PK 1234 was greater when co-transfected with yeast tRNA than when in competition with *wt* mCherry replicon, the ptGFP signal was reduced at passage two and was at background level by passage three (figure 5C). Replication of the mCherry *wt* replicon was not influenced by co-transfection with the ptGFP constructs (figure 5D), as expected. Together these data suggest that the minor replicative advantage conferred by multiple PKs are quickly compounded over multiple replication cycles to provide a replicative advantage.

### Presence of a PK is essential for the production of infectious virus

As replicons lacking all PKs could replicate and replicons with reduced numbers of PKs appeared to be at a competitive disadvantage compared to the *wt* construct, we investigated the consequences of PK manipulation on the complete viral life cycle. The ∆PK 34, ∆PK 234 and C11 ∆PK 1234 mutations were introduced into an FMDV infectious clone by replacement of sequence encoding ptGFP with that encoding the O1K structural proteins. RNA transcripts were transfected into BHK-21 cells alongside a *wt* O1K viral transcript and blind passaged 5 times by transferring the cell supernatant at 24 hours post transfection onto naïve BHK-21 cells. After 5 blind passages, recovered virus was harvested and sequenced to check for compensatory or reversion mutations.

*Wt*, C11, ∆PK 34 and ∆PK 234 constructs all resulted in the production of infectious virus as was expected from the replicon experiments, with no alteration to input sequence. However, the C11 ∆PK 1234, which replicated (albeit to a lesser degree) as a replicon, produced no recoverable infectious virus (Table 1). Interestingly, there were differences noted in both the rate of CPE (figure 6A) and plaque size (figure 6B-C) of ∆PK 34 and ∆PK 234 when compared to the *wt* O1K virus. Rate of CPE was monitored by infecting BHK-21 cells with a known MOI (0.01) of recovered virus, cells were then monitored for signs of CPE (shown as a decrease in cell confluency) as measured by an automated imaging platform (Incucyte Zoom). Both ∆PK 34 and ∆PK 234 showed delayed onset of CPE with ∆PK 34 being the slowest, initial CPE occurring at approximately 39 hours and 29 hours post infection respectively, compared to the 22 hours seen in the *wt* control. This mirrored plaque assay data where ∆PK 34 displayed a significantly smaller plaque phenotype when compared to the *wt* control (average of 13.8 pixels compared to 37.4), the slower rate of CPE seen in ∆PK 234 made a small, but not significant difference (average 31.9 pixels).

**Table 1.** Virus could not be recovered when all PKs are deleted. FMDV infectious clone RNA was transfected into BHK-21 cells and appearance of CPE observed over 4 sequential passages. At the fifth passage virus was harvested and sequenced to observe for any changes within the sequence. Presence of CPE indicated with ‘Y’ while ‘N’ represent no CPE seen.

|  | Appearance of CPE |  |  |  |  |
| --- | --- | --- | --- | --- | --- |
| Recovered Virus | Passage 1 | Passage 2 | Passage 3 | Passage 4 | Sequence of Rescued Virus |
| WT | Y | Y | Y | Y | No Change |
| C11 | Y | Y | Y | Y | No Change |
| ΔPK34 | Y | Y | Y | Y | No Change |
| ΔPK234 | Y | Y | Y | Y | No Change |
| ΔPK1234 | N | N | N | N | N/A |

**Figure 6.**
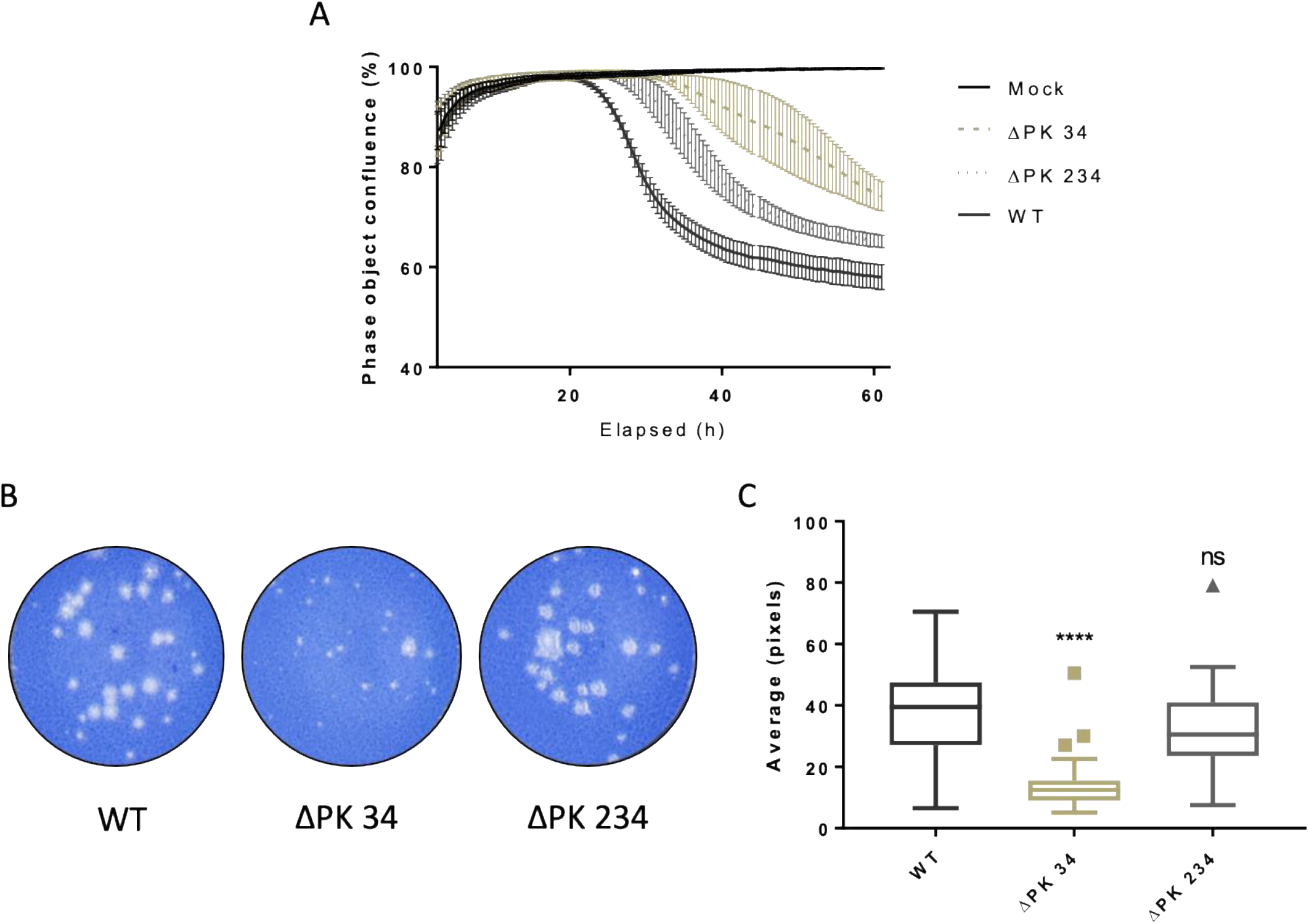
Recovered virus showed a delay in rate of CPE and a small plaque phenotype. BHK-21 cells were infected with *wt*, ΔPK 34 and ΔPK 234 virus at a MOI of 0.01, alongside a mock infected control and cell confluency (shown as phase object confluence %) monitored every half an hour for 62 hours using an Incucyte Zoom (A). Representative plaque assay of BHK-21 cells infected with *wt*, ΔPK 34 and ΔPK 234 viruses. Recovered virus from passage 5 of blind passaging was infected onto BHK-21 cells, cells were fixed and stained 48 hours post infection (B). Virus plaques were imaged and size of plaques measured using Image J, all plaques per well were counted, additional wells were used until a minimum plaque count of 40 was reached. Significance is shown when compared to the *wt*, box and whisker plots were made using the Tukey method.

Since C11 ∆PK 1234 produced no infectious virus the ability of the full-length genome lacking PKs to replicate was investigated. BHK-21 cells were transfected with the same RNA transcripts as above alongside additional controls, mock-transfected and transfected with *wt* and treated with 3 mM GuHCl (a replication inhibitor) as negative controls. Six hours post-transfection, cells were harvested, fixed and labelled with an anti-3A antibody and fluorescent secondary antibody. Cells were then analysed using flow cytometry and anti-3A antibody signal used as an indirect measure of genome replication (figure 7). The results were similar to those of the replicon experiments and showed that all the modified virus genomes were able to undergo robust replication. The inability of the C11 ∆PK 1234 genome to support production of infectious virus despite being able to replicate after transfection into cells, is consistent with a requirement for RNA structure within the PK region being required for virus assembly.

**Figure 7.**
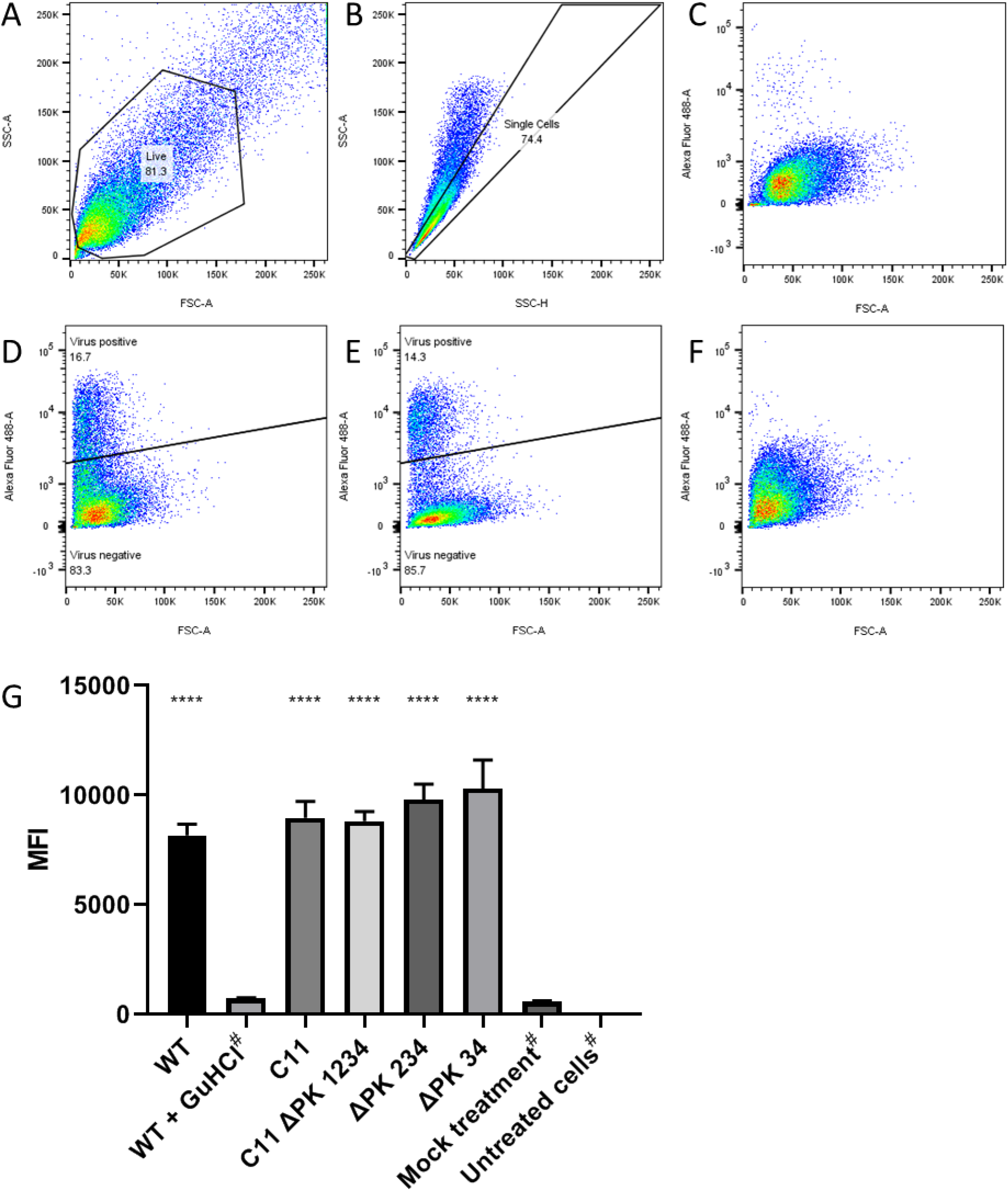
PKs are not essential for viral genome replication. BHK-21 cells were transfected with RNA transcribed from infectious clones of *wt*, ΔPK 34, ΔPK 234 and C11 ΔPK 1234. Non-transfected cells and a *wt* transfection treated with 3 mM GuHCl were used as negative controls. Cells were harvested, fixed and labelled using an anti-3A antibody and fluorescent anti-mouse secondary before separation by flow cytometry. Gates were used to select for live cell and single cell populations, and virus positive/negative populations were identified based on levels of Alexa-488 fluorescence. Where distinct virus positive and negative populations exist, gates were drawn to separate these in order to determine the mean fluorescent intensity (MFI) of the virus positive cells. Where no clear separate populations exist (*wt* with GuHCl, mock treatment and untreated cells) gates could not be drawn and therefore the total MFI has been reported (denoted by #). Representative images have been shown here for the live cell gate from the *wt* virus transfection (A), the single cell gate from the *wt* virus transfection (B) and the relative fluorescence of the cells with mock antibody treatment (C), *wt* virus (D), C11 ΔPK 1234 virus (E) and *wt* virus transfection with 3 mM GuHCl (F). The experiment was performed in triplicate and the MFI values for each condition were calculated (G). The error bars represent the SEM. Significance is shown compared to the *wt* plus 3 mM GuHCl control (**** P < 0.0001).

## Discussion

The 5′ UTR of FMDV is unique amongst picornaviruses due to its large size and the presence of multiple RNA elements, some of which still have unknown function. One of these features is a series of repeated PKs varying in number from 2-4, depending on virus strain. In this study, we sequentially deleted or mutated the PKs to help understand their role in the viral life cycle. We also confirmed the predicted PK structures by SHAPE mapping, although there may be fewer strong interactions maintaining the PKs than was previously predicted. This may indicate high conformational flexibility of this region of the genome. SHAPE mapping was also supported by mutation of predicted key interactions between nucleotides in the loop and downstream, disruption of which reduced replication to that of the C11 ∆PK 1234 replicon.

Although all viruses isolated to date contain at least two PKs, replicons or viruses containing a single PK were still replication competent. However, replicons with more than a single PK were found to have a competitive advantage over replicons with a single PK when sequentially passaged. Replicons lacking all PKs displayed poor passaging potential even when co-transfected with yeast tRNA, reinforcing the observation of a significant impact in replication. Moreover, viruses recovered from genomes with reduced numbers of PKs were slower growing and produced smaller plaques. In addition, these differences were more pronounced in more relevant cells lines (i.e. in MDBK cells compared to BHK 21 cells). It is likely that each of the PKs is functionally competent as no differences was seen between replicons congaing a single copy of PK1 or PK4. This observation is consistent with a previous report of deletion of PK1, along with the Poly-C-tract, with no adverse effect in viral replication (37). This also supports our findings that the truncation of the Poly-C-tract to create the C11 construct had no effect on replicon replication in the cell lines tested. As has been described with Mengo virus, it is possible that the role of the poly-C-tract is essential in other aspects of the viral lifecycle which cannot be recapitulated in a standard tissue culture system (39).

The presence of at least two PKs in all viral isolates sequenced so far suggests that multiple PKs confer a competitive advantage in replication. Here we showed by sequential passage that replicons containing at least two PKs were maintained at a level similar to *wt*, but replicons containing only one PK showed a persistent decline. It is unclear why some viral isolates contain two, three or four PKs is still unknown, but this may be stochastic variation or may reflect subtle effects of host range or geographical localisation.

Surprisingly, although removal of all four PKs resulted in a significant decrease in replicon and viral genome replication, replication was not abolished, showing that PKs are not essential to support genome replication. However, deletion of all PKs from an infectious clone completely abolished the ability to recover infectious virus. This suggests that the genome lacking all PKs is defective in a function associated with virion assembly and is compatible with our evidence for the presence of a packaging signal in a similar location on the genome to PK1 (22). It is possible that structural flexibility at this site in the genome allows the RNA to adopt alternate conformations with different roles in genome replication and virion assembly. A functional requirement for multiple RNA conformations may explain the relatively weak interactions between nucleotides involved in stabilising the PK motif as observed by SHAPE analysis or by structural prediction.

## Acknowledgments

This work was supported by funding from the Biotechnology and Biological Sciences Research Council (BBSRC) of the United Kingdom (research grant BB/K003801/1). Additionally, the Pirbright Institute receives grant-aided support from the BBSRC (projects BB/E/I/00007035, BB/E/I/00007036 and BBS/E/I/00007037).

## References

1. Knight-Jones TJD, Rushton J. 2013. The economic impacts of foot and mouth disease – What are they, how big are they and where do they occur? Prev Vet Med 112:161–173.

2. Mahapatra M, Parida S. 2018. Foot and mouth disease vaccine strain selection: current approaches and future perspectives. Expert Rev Vaccines 17:577–591.

3. Park J-H. 2013. Requirements for improved vaccines against foot-and-mouth disease epidemics. Clin Exp Vaccine Res 2:8–18.

4. Stenfeldt C, Eschbaumer M, Rekant SI, Pacheco JM, Smoliga GR, Hartwig EJ, Rodriguez LL, Arzt J. 2016. The Foot-and-Mouth Disease Carrier State Divergence in Cattle. J Virol 90:6344–64.

5. Carrillo C, Tulman ER, Delhon G, Lu Z, Carreno A, Vagnozzi A, Kutish GF, Rock DL. 2005. Comparative genomics of foot-and-mouth disease virus. J Virol 79:6487–504.

6. Curry S, Fry E, Blakemore W, Abu-Ghazaleh R, Jackson T, King A, Lea S, Newman J, Stuart D. 1997. Dissecting the roles of VP0 cleavage and RNA packaging in picornavirus capsid stabilization: the structure of empty capsids of foot-and-mouth disease virus. J Virol 71:9743–52.

7. Gao Y, Sun S-Q, Guo H-C. 2016. Biological function of Foot-and-mouth disease virus non-structural proteins and non-coding elements. Virol J 13:107.

8. Herod MR, Gold S, LaseckaDykes L, Wright C, Ward JC, McLean TC, Forrest S, Jackson T, Tuthill TJ, Rowlands DJ, Stonehouse NJ. 2017. Genetic economy in picornaviruses: Foot-and-mouth disease virus replication exploits alternative precursor cleavage pathway. PLOS Pathog 13:e1006666.

9. Tulloch F, Pathania U, Luke GA, Nicholson J, Stonehouse NJ, Rowlands DJ, Jackson T, Tuthill T, Haas J, Lamond AI, Ryan MD. 2014. FMDV replicons encoding green fluorescent protein are replication competent. J Virol Methods 209:35–40.

10. Herod MR, Tulloch F, Loundras E-A, Ward JC, Rowlands DJ, Stonehouse NJ. 2015. Employing transposon mutagenesis to investigate foot-and-mouth disease virus replication. J Gen Virol 96:3507–3518.

11. Mellor EJC, Brown F, Harris TJR. 1985. Analysis of the Secondary Structure of the Poly(C) Tract in Foot-and-Mouth Disease Virus RNAs. J Gen Virol 66:1919–1929.

12. Clarke BE, Brown AL, Currey KM, Newton SE, Rowlands DJ, Carroll AR. 1987. Potential secondary and tertiary structure in the genomic RNA of foot and mouth disease virus. Nucleic Acids Res 15:7067–7079.

13. Nayak A, Goodfellow IG, Woolaway KE, Birtley J, Curry S, Belsham GJ. 2006. Role of RNA structure and RNA binding activity of foot-and-mouth disease virus 3C protein in VPg uridylylation and virus replication. J Virol 80:9865–75.

14. Belsham GJ, Brangwyn JK. 1990. A region of the 5’ noncoding region of foot-and-mouth disease virus RNA directs efficient internal initiation of protein synthesis within cells: involvement with the role of L protease in translational control. J Virol 64:5389–95.

15. Yang F, Zhu Z, Cao W, Liu H, Wei T, Zheng M, Zhang K, Jin Y, He J, Guo J, Liu X, Zheng H. 2020. Genetic Determinants of Altered Virulence of Type O Foot-and-Mouth Disease Virus. J Virol.

16. Kloc A, Diaz-San Segundo F, Schafer EA, Rai DK, Kenney M, de los Santos T, Rieder E. 2017. Foot-and-mouth disease virus 5’-terminal S fragment is required for replication and modulation of the innate immune response in host cells. Virology 512:132–143.

17. Kloc A, Rai DK, Rieder E. 2018. The roles of picornavirus untranslated regions in infection and innate immunity. Front Microbiol. Frontiers Media S.A.

18. Zhu Z, Yang F, Cao W, Liu H, Zhang K, Tian H, Dang W, He J, Guo J, Liu X, Zheng H. 2019. The Pseudoknot Region of the 5’ Untranslated Region Is a Determinant of Viral Tropism and Virulence of Foot-and-Mouth Disease Virus. J Virol 93.

19. Mohapatra JK, Pawar SS, Tosh C, Subramaniam S, Palsamy R, Sanyal A, Hemadri D, Pattnaik B. 2011. Genetic characterization of vaccine and field strains of serotype A foot-andmouth disease virus from India. Acta Virol 55:349–352.

20. Escarmís C, Dopazo J, Dávila M, Palma EL, Domingo E. 1995. Large deletions in the 5’-untranslated region of foot-and-mouth disease virus of serotype C. Virus Res 35:155–67.

21. Carocci M, Bakkali-Kassimi L. 2012. The encephalomyocarditis virus. Virulence 3:351–67.

22. Wutz G, Auer H, Nowotny N, Grosse B, Skern T, Kuechler E. 1996. Equine rhinovirus serotypes 1 and 2: relationship to each other and to aphthoviruses and cardioviruses. J Gen Virol 77:1719–1730.

23. Chapman EG, Moon SL, Wilusz J, Kieft JS. 2014. RNA structures that resist degradation by Xrn1 produce a pathogenic Dengue virus RNA. Elife 3.

24. Kieft JS, Rabe JL, Chapman EG. 2015. New hypotheses derived from the structure of a flaviviral Xrn1-resistant RNA: Conservation, folding, and host adaptation. RNA Biol 12:1169–77.

25. Gultyaev AP, Olsthoorn RCL. 2010. A family of non-classical pseudoknots in influenza A and B viruses. RNA Biol 7:125–9.

26. Moss WN, Dela-Moss LI, Priore SF, Turner DH. 2012. The influenza A segment 7 mRNA 3’ splice site pseudoknot/hairpin family. RNA Biol 9:1305–10.

27. Plant EP, Dinman JD. 2008. The role of programmed-1 ribosomal frameshifting in coronavirus propagation. Front Biosci 13:4873–81.

28. Herod MR, Ferrer-Orta C, Loundras E-A, Ward JC, Verdaguer N, Rowlands DJ, Stonehouse NJ. 2016. Both cis and trans Activities of Foot-and-Mouth Disease Virus 3D Polymerase Are Essential for Viral RNA Replication. J Virol 90:6864–6883.

29. Karabiber F, McGinnis JL, Favorov O V., Weeks KM. 2013. QuShape: Rapid, accurate, and best-practices quantification of nucleic acid probing information, resolved by capillary electrophoresis. RNA 19:63–73.

30. Darty K, Denise A, Ponty Y. 2009. VARNA: Interactive drawing and editing of the RNA secondary structure. Bioinformatics 25:1974–1975.

31. King AMQ, Blakemore WE, Ellard FM, Drew J, Stuart DI. 1999. Evidence for the role of His-142 of protein 1C in the acid-induced disassembly of foot-and-mouth disease virus capsids. J Gen Virol 80:1911–1918.

32. Herod MR, Gold S, Lasecka-Dykes L, Wright C, Ward JC, McLean TC, Forrest S, Jackson T, Tuthill TJ, Rowlands DJ, Stonehouse NJ. 2017. Genetic economy in picornaviruses: Foot-and-mouth disease virus replication exploits alternative precursor cleavage pathways. PLOS Pathog 13:e1006666.

33. Logan G, Freimanis GL, King DJ, Valdazo-González B, Bachanek-Bankowska K, Sanderson ND, Knowles NJ, King DP, Cottam EM. 2014. A universal protocol to generate consensus level genome sequences for foot-and-mouth disease virus and other positive-sense polyadenylated RNA viruses using the Illumina MiSeq. BMC Genomics 15:828.

34. Acevedo A, Andino R. 2014. Library preparation for highly accurate population sequencing of RNA viruses. Nat Protoc 9:1760–1769.

35. Peng Y, Leung HCM, Yiu SM, Chin FYL. 2012. IDBA-UD: a de novo assembler for single-cell and metagenomic sequencing data with highly uneven depth. Bioinformatics 28:1420–1428.

36. Altschul SF, Gish W, Miller W, Myers EW, Lipman DJ. 1990. Basic local alignment search tool. J Mol Biol 215:403–410.

37. Rieder E, Bunch T, Brown F, Mason PW. 1993. Genetically engineered foot-and-mouth disease viruses with poly(C) tracts of two nucleotides are virulent in mice. J Virol 67:5139–45.

38. Herod MR, Ferrer-Orta C, Loundras E-A, Ward JC, Verdaguer N, Rowlands DJ, Stonehouse NJ. 2016. Both cis and trans Activities of Foot-and-Mouth Disease Virus 3D Polymerase Are Essential for Viral RNA Replication. J Virol 90:6864–6883.

39. Martin L, Duke G, Osorio J, Hall D, Palmenberg A. 1996. Mutational analysis of the mengovirus poly(C) tract and surrounding heteropolymeric sequences. J Virol 70:2027–2031.

